# Phylogenetic diversity and activity screening of cultivable actinobacteria isolated from marine sponges and associated environments from the western coast of India

**DOI:** 10.1101/2020.01.09.901108

**Authors:** Ulfat Baig, Neelesh Dahanukar, Neha Shintre, Ketki Holkar, Anagha Pund, Uttara Lele, Tejal Gujarathi, Kajal Patel, Avantika Jakati, Ruby Singh, Harshada Vidwans, Vaijayanti Tamhane, Neelima Deshpande, Milind Watve

**Author notes:** Correspondence and requests for materials should be addressed to N. Dahanukar.

## Abstract

Phylogenetic diversity of cultivable actinobacteria isolated from sponges (*Haliclona* spp.) and associated environments of intertidal zones, along the northern parts of west coast of India, were studied using 16S rRNA gene sequences. A subset of actinobacteria were screened for three activities, namely predatory behavior, antibacterial activity and enzyme inhibition. We recovered 237 isolates of actinobacteria belonging to 19 families and 28 genera, which could be attributed to 95 putative species using maximum likelihood partition and 100 putative species using Bayesian partition in Poisson Tree Processes. Although the trends in the discovery of actinobacterial genera isolated from sponges was consistent with previous studies from different study areas, we provide first report of nine actinobacterial species from sponges. We observed widespread non-obligate epibiotic predatory behavior in eight actinobacterial genera and we provide first report of predatory activity in *Brevibacterium*, *Glutamicibacter*, *Micromonospora*, *Nocardiopsis*, *Rhodococcus* and *Rothia*. Sponge associated actinobacteria showed significantly more predatory behavior than environmental isolates. While antibacterial activity by actinobacterial isolates mainly affected Gram-positive target bacteria with little to no effect on Gram-negative bacteria, predation targeted both Gram-positive and Gram-negative prey with equal propensity. Actinobacterial isolates from both sponge and associated environment produced inhibitors of serine proteases and angiotensin converting enzyme. Predatory behavior was strongly associated with inhibition of trypsin and chymotrypsin. Our study suggests that sponge and associated environment of western coast of India are rich in actinobacterial diversity with widespread predatory activity, antibacterial activity and production of enzyme inhibitors. Understanding diversity and associations among various actinobacterial activities, with each other and the source of isolation, can provide new insights in marine microbial ecology and provide opportunities to isolate novel therapeutic agents.

## INTRODUCTION

The marine ecosystem is not only diverse with respect to microorganisms found in it but also the natural products being synthesized by these microorganisms (Ward and Bora, 2006; Taylor et al., 2007; Lam, 2006). Actinobacteria are among the taxa rich in secondary metabolites (Barka et al., 2016) and are widely distributed in diverse habitats including soil, marine and freshwaters and sediments (Ward and Bora, 2006; Taylor et al., 2007; Tan et al., 2015; Brasel et al., 2019; Mincer et al., 2002; Kokare et al., 2004). They are also not uncommon in extreme environments (Jose and Jebakumar, 2014; Pathom-Aree et al., 2006; Mohammadipanah and Wink, 2016; Shivlata and Tulasi, 2015; Riquelme et al., 2015; Yang et al., 2015) and are also found as endobiotic symbionts of higher organisms (Taylor et al., 2007; Li et al., 2015; Mahmoud and Kalendar, 2016; Trujillo et al., 2015). They belong to the phylum Actinobacteria and represent one of the major phyla within the bacterial domain (Goodfellow, 2015). They are aerobic, spore forming, Gram-positive bacteria, which often produce diffusible pigments, and occur as cocci or rods, branched filaments, aerial or substrate mycelium (Goodfellow, 2015). The marine ecosystems are believed to have a wide range of unexplored diversity of actinobacteria (Montalvo et al., 2005) and their metabolites (Taylor et al., 2007; Lam, 2006; Manivasagan et al., 2005) with diverse biological activities like anticancer (Olano et al., 2009), anti-inflammatory (Trischman et al., 1994), antibiotic (Pimentel-Elardo et al., 2010; Cheng et al., 2015; Gandhimathi et al., 2008), cytotoxic (Abdelfattah et al., 2016) and enzyme inhibitory (Manivasagan et al., 2015; Imada, 2005) activity. Watve et al. (2001) estimated that the genus *Streptomyces* alone is capable of producing up to 10^5^ different metabolites, majority of which remain unexplored. Of 23,000 medicinally important metabolites produced by marine microorganisms 70% are contributed by actinobacteria (Mahapatra et al., in press). Till date, eight genera of actinobacteria have been reported to produce secondary metabolites and 267 products have been reported from 96 marine actinobacteria (Subramani and Sipkema, 2019)^31^.

Ecologically it is difficult to understand the production of extracellular metabolites or enzymes by aquatic bacteria, since any molecule secreted outside the cell can be quickly washed off. Extracellular products could be useful to the producer only in viscous or partially enclosed environments. In the marine environment, sponges are likely to provide such closed environment for bacteria. Sponges are filter feeders and collect small nutrient particles including bacteria. This makes the environment locally nutrient rich in an otherwise oligotrophic surroundings. Bacteria, especially actinobacteria, isolated from these sponges may live in a symbiotic relationship that helps the host in defense against predation, sponge skeleton stabilization, translocation of metabolites and help in nutritional process (Taylor et al., 2007; Li et al., 2015; Montalvo et al., 2005; Pimentel-Elardo et al., 2010; Cheng et al., 2015; Gandhimathi et al., 2008; Lee et al., 2009; Thomas et al., 2010). In addition, since sponges are sessile and lack other anti-predator defenses, secondary metabolites of bacteria can provide them with chemical defense (Lee et al., 2001). Therefore, we expect more secondary metabolite related activities from sponge-associated actinobacteria.

Sponge-associated actinobacteria are likely to have another ecological role. Among actinobacteria at least three genera, namely *Agromyces, Streptomyces* and *Streptoverticillium*, are shown to be predators that kill and feed on other live bacterial cells (Casida, 1980; 1983; Kumbhar et al., 2014). Kumbhar and Watve (2013) argued that antibiotic activity might have evolved primarily as a weapon in predation. However, the expression of secondary metabolites during predation may be independent of antibiotic expression in pure culture; the latter is likely to have evolved for mutualism with higher animal or plant hosts (Harir et al., 2018; Van der Meij et al., 2017). Further, for a niche of predation in association with sponge, the predatory species needs to protect itself from the digestive enzymes of the sponge as well as its own enzymes used for predation. Therefore, predatory actinobacteria are also expected to have efficient inhibitors of lytic enzymes.

In this study, we prepared an inventory of cultivable actinobacteria from sponges and associated environments of intertidal zones along the northern parts of west coast of India and studied their molecular diversity based on 16S rRNA gene sequences. We screened a subset of randomly selected cultures for predatory activity, antibiotic production and enzyme inhibition and tested their associations with each other and with the isolation source to test the hypotheses mentioned earlier.

## MATERIALS AND METHODS

### Sample collection

Small tissue samples (less than one gram) of marine sponges (*Haliclona* spp.) were collected at the time of low tide along Maharashtra and Goa coast (18–15°N and 73–74°E) of India during April 2014 to October 2018 without damaging the sponge or its associated environment. Specimens were rinsed and flushed with sterile media to remove debris and loosely attached microbes. Each sponge sample was collected in labeled polystyrene tubes with lids containing sterile Poor Ravan Saline (Watve et al., 2000) and ZoBell Marine broth (ZoBell, 1941). Sediment, water and air samples were collected from the same environment as that of the sponge and were collectively considered as environmental samples. The samples were brought to laboratory maintaining cold chain and were immediately processed for microbial culturing.

### Isolation and maintenance of cultivable actinobacteria

Each sample was subjected to pre-heat treatment at 60°C for 15 minutes to eliminate non-sporulating bacteria. Sponge tissue (0.1 cm^3^) was homogenized in sterile medium and vortexed for 5 minutes. Tubes were left undisturbed for two minutes. From the resulting supernatant serial 10 fold dilutions upto 10^−5^ were made and 0.1 ml sample was spread into triplicates on petri plates containing sterile medium. We used two media the Zobell Marine Agar (ZMA) and Poor Ravan Saline Agar (PRSA) with and without antibiotic chloramphenicol (25 μg/ml). Plates were incubated at 30°C for 7 days in the case of ZMA and 21 days for PRSA. Plates were observed regularly for the growth of actinobacteria. Bacterial colonies that showed resemblance to actinobacteria under light microscope were purified several times on the respective media. In all 237 actinobacterial isolates were selected and were re-streaked for making pure cultures. Colonies were labeled as per Maharashtra Gene Bank (MGB) project code and preserved on ZMA slants at 4°C for further use. Similarly, glycerol (18%) stocks were prepared and maintained at −20°C for long term storage. Actinobacterial cultures are deposited in the Microbial Culture Collection (MCC) of National Centre for Microbial Resource, National Center for Cell Sciences, Pune, India (accession numbers are provided in the Supplementary Table S1).

### Genetic identification, phylogeny and species delimitation

Actinobacterial isolates were outsourced for near complete 16S rRNA gene sequencing. Gene sequences used for the study are deposited in the GenBank database under the accession numbers MN339687–MN339897 and MT598037–MT598065 (Supplementary Table S1)). Sequences were checked in BLAST (Altschul et al., 1990) to find the closest sequences available in the GenBank database (href="http://www.ncbi.nlm.nih.gov). Four species of Firmicutes, namely *Bacillus paralicheniformis* (MCC 6306), *B. thuringiensis* (MCC 7835), *B. subtilis* (MCC 6386) and *B. halotolerans* (MCC 8381), were used as outgroups (GenBank accession numbers MN339894–MN339897 respectively).

Gene sequences were aligned using MUSCLE (Edgar, 2004) implemented in MEGA 7 (Kumar et al., 2016). Final aligned matrix had 1595 sites. Best nucleotide substitution model was determined using ModelFinder (Kalyaanamoorthy et al., 2017) based on Bayesian information criterion (Schwarz, 1978; Nei and Kumar, 2000). Maximum likelihood analysis was performed in IQ-TREE (Nguyen et al., 2015) with ultrafast bootstrap support (Hoang et al., 2018) for 1000 iterations. Phylogenetic tree was edited in FigTree v1.4.2 (Rambaut, 2009).

To understand putative number of actinobacterial species we performed species delimitation based on Poisson Tree Processes (Zhang et al., 2013) with maximum likelihood partitioning (mPTP) and Bayesian partitioning (bPTP). Maximum likelihood tree was used to delimit species by setting the parameter values as follows: MCMC generations = 100,000, Thinning = 100, Burn-in = 0.1 and seed = 123.

We have identified all isolates up to genus level, while operational taxonomic units, in terms of putative species, are provided based on mPTP and bPTP methods (see Supplementary Table S1). Only in the text, some isolates are assigned to known species based on BLAST search and sequence identity more than 99%.

### Screening for activities

Out of 237 actinobacterial isolates, 50 isolates were randomly selected for screening of three activities, namely predation, antibiotic production and production of enzyme inhibition.

### Target bacteria used for predation and antibiotic screening

Test bacteria, used for checking actinobacterial predation and antibiotic production, were obtained from MCC and National Collection of Industrial Microorganisms (NCIM), National Chemistry Laboratory, Pune, India. Fourteen bacteria, namely *Acetobacter pasterianus* (NCIM 2317), *Alcaligenes fecalis* (NCIM 2262), *Bacillus subtilis* (NCIM 2063), *Enterobacter fecalis*, *Escherichia coli* (NCIM 2184), *Klebsiella pneumonae* (NCIM 2957), *Micrococcus luteus* (NCIM 2673), *Mycobacterium smegmatis* (NCIM 5138), *Proteus vulgaris* (NCIM 2172), *Pseudominas aeruginosa* (NCIM 5029), *Salinicoccus roseus* (MCC 7574), *Salmonella enterica* (NCIM 2501), *Serretia marcescens* (NCIM 2919) and *Staphylococcus aureus* (NCIM 2121), were used as target species for screening.

### Screening for actinobacterial predatory growth

Growth of predator with the zone of clearance on prey cells was considered as predation as defined earlier (Kumbhar et al., 2014). The method for the preparation of prey cells was modified from Kumbhar et al. (2014). Pure cultures of the prey species were inoculated on nutrient agar plates to check the purity and were later re-inoculated in nutrient broth. Inoculated flasks were incubated at 37°C for 24 h. Broth was centrifuged at 7000 rpm for 10 minutes to concentrate cells using Eppendorf centrifuge 5810R. Cells were washed thrice with sterile distilled water to remove traces of nutrient broth. Pellet was suspended in saline to obtain a thick suspension of optical density of 1.0 at 600 nm. Lawn of prey cells was spread on water agarose plate and plates were incubated at 37°C for 40 minutes. Actinobacterial culture was spot inoculated on pre incubated plates. These plates were incubated at room temperature for 48–72 h at 30°C. Plates with plaque were examined visually and by using 4x and 45x magnification under light microscope. Prey and predator control plates were used for comparison. Each experiment consisted of triplicate sets of plates, as well as one predator control for testing growth of actinobacterial predator without prey. In addition, there was a prey control to demonstrate viable and independent growth of prey without predator. In either controls there was no zone of clearance indicating there was no predation in the presence of predator or prey alone.

### Screening for antibacterial activity using conventional cross streak method

Selected actinobacterial cultures were screened for antibacterial activity by cross streak method (Velho-Pereira and Kamat, 2011; Valli et al., 2012). Test organism was streaked as a straight line along the diagonal of the petri dish with sterile ZMA medium. The isolated pure colony of actinobacteria was inoculated as a single streak perpendicular to the central streak. Streaking was done from the edge of the plate to the test organism growth line. Plates were incubated at 37°C for 18 h. The microbial inhibition was observed by determining zone of clearance around the sensitive organisms. Control plates of the same medium with the streak of test bacteria and without the streak of actinobacteria growth was used to observe the normal growth of the test bacteria.

### Screening for enzyme inhibitors

Actinobacterial cultures were screened for their ability to inhibit the activity of serine proteases and angiotensin converting enzyme (ACE). Three different serine proteases i.e., Subtilisin, Trypsin and α-Chymotrypsin were used for screening of inhibitory activity. Protease inhibitor activity was studied using unprocessed X-ray films and spot-test method (Cheung, 1991) with modifications. As described by Tripathi et al. (Tripathi et al., 2011), dilutions of pure enzyme were first spotted on gelatine coated films. Lowest dilution showing complete clearance (indicating complete digestion of gelatine) was chosen for further studies. Pure enzyme (100 μg/ml) was incubated with equal quantity of cell free supernatant of actinobacterial isolates for 10 minutes and transferred to untreated X-ray-Fuji Medical X-ray, HRU grade-films. The mixtures were allowed to react for 15 minutes at room temperature and results were recorded after washing the x-ray films under running water. Unprocessed X-ray films contain a layer of gelatine on their surface, which acts as a substrate for various proteolytic enzymes. Degradation of gelatine gives a clear zone at the site of activity. Thus, upon action of the proteases, clear zones were seen on unprocessed x-ray films, at the site of inoculation, whereas, if the gelatine layer remains intact, no clearance is observed. No clearance on the films indicated presence of protease inhibitors.

ACE acts on a specific substrate N-Hyppuryl-His-Leu (HHL) to liberate hippuric acid and His-Leu. Liberated hippuric acid was detected spectrophotometrically. Upon reaction of the enzyme with ACE inhibitors, the enzyme becomes inactive and this is measured in terms of lower levels of hippuric acid released. Protocol suggested by Cushman and Cheung (1971) was used with certain modifications and hippuric acid liberated was checked using method suggested by Ng et al. (2008). Equal amount of ACE and cell free supernatants (10 μl each) were allowed to react at 37°C. After 10 minutes 20 μl of HHL was added to the reaction mixture and reaction was continued for 30 mins at 37°C. The reaction was stopped by addition of 40 μl of 1 N HCl. Blank was prepared by addition of HCl before addition of the substrate. Positive enzyme control was prepared by incubating enzyme with un-inoculated broth. Liberated hippuric acid was extracted in 90 μl ethyl acetate by vigorous shaking. Ethyl acetate layer was collected in a fresh vial and allowed to dry in water bath of 50°C. The liberated hippuric acid was diluted in 150 μl distilled water and absorbance was checked at 228 nm. Zero was adjusted using distilled water. Test vials with more than 15% inhibition of ACE were considered as positive for ACE inhibitor.

## RESULTS

### Actinobacterial phylogenetic diversity in sponge and associated environment

Actinobacteria from sponges and associated environments showed a rich phylogenetic diversity (Figure 1). We obtained 237 actinobacterial isolates, from sponge and associated environments, belonging to 19 families and 28 genera (Supplementary Table S1). Species delimitation based on mPTP suggested that these isolates belong to 95 putative species, while bPTP suggested 100 putative species. The two species delimitation methods, mPTP and bPTP, differed in the groups of species under genera *Micrococcus*, *Rhodococcus* and *Streptomyces* (Supplementary Table S1). Air was generally devoid of actinobacteria and we recovered only three isolates from air, belonging to genera *Brachybacterium*, *Brevibacterium* and *Rhodococcus*, as compared to 39 isolates from water, 105 isolates from sediment and 90 isolates from sponge.

**Figure 1.**
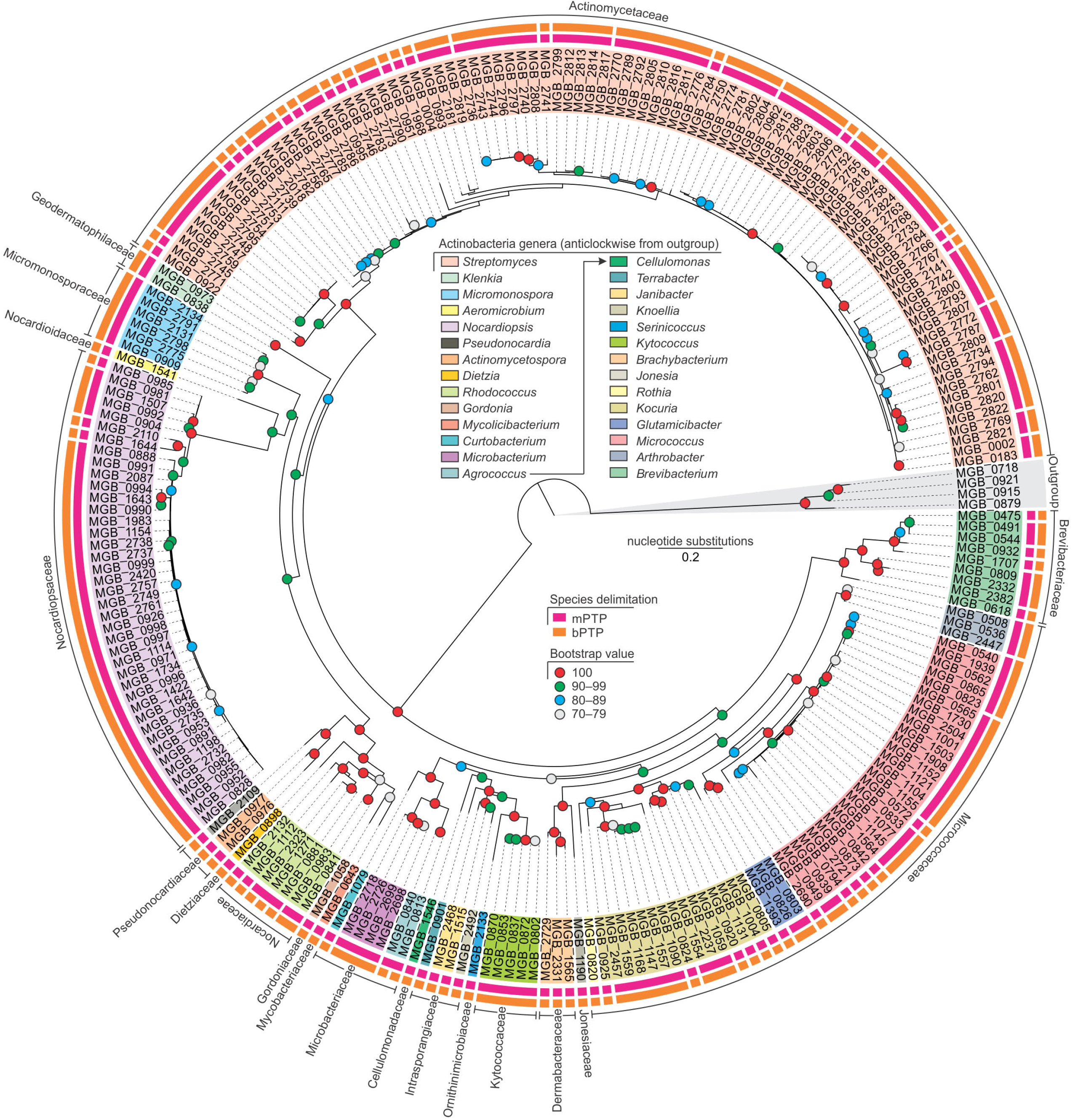
Maximum likelihood phylogenetic tree of actinobacterial isolates based on TIM3+F+I+G4 nucleotide substitution model (lnL of consensus tree: −18684.58). Firmicutes belonging to genus *Bacillus* were used as outgroups.

From sponges, 18 genera under 14 families belonging to 56 putative species (Table 1). From the sponge-associated environment, 22 genera under 15 families were recorded belonging to 64 putative species as per mPTP and 65 putative species as per bPTP. A total of 12 genera under 9 families and 28 putative species based on mPTP and 25 putative species based on bPTP were common to both sponge and associated environment.

**Table 1.** Putative number of species of actinobacterial genera based on PTP and bPTP methods isolated from sponge, associate environment and both sources.

| Family | Genus | Sponge |  | Environment |  | Both |  |
| --- | --- | --- | --- | --- | --- | --- | --- |
|  |  | mPTP | bPTP | mPTP | bPTP | mPTP | bPTP |
| Actinomycetaceae | *Streptomyces* | 23 | 23 | 24 | 25 | 12 | 11 |
| Brevibacteriaceae | *Brevibacterium* | 5 | 5 | 2 | 2 | 1 | 1 |
| Cellulomonadaceae | *Cellulomonas* | 0 | 0 | 1 | 1 | 0 | 0 |
| Dermabacteraceae | *Brachybacterium* | 1 | 1 | 2 | 2 | 0 | 0 |
| Dietziaceae | *Dietzia* | 0 | 0 | 1 | 1 | 0 | 0 |
| Geodermatophilaceae | *Klenkia* | 1 | 1 | 1 | 1 | 1 | 1 |
| Gordoniaceae | *Gordonia* | 1 | 1 | 0 | 0 | 0 | 0 |
| Intrasporangiaceae | *Janibacter* | 0 | 0 | 2 | 2 | 0 | 0 |
|  | *Knoellia* | 0 | 0 | 1 | 1 | 0 | 0 |
|  | *Terrabacter* | 0 | 0 | 1 | 1 | 0 | 0 |
| Jonesiaceae | *Jonesia* | 1 | 1 | 0 | 0 | 0 | 0 |
| Kytococcaceae | *Kytococcus* | 0 | 0 | 1 | 1 | 0 | 0 |
| Microbacteriaceae | *Agrococcus* | 1 | 1 | 1 | 1 | 1 | 0 |
|  | *Curtobacterium* | 0 | 0 | 1 | 1 | 0 | 0 |
|  | *Microbacterium* | 1 | 1 | 1 | 1 | 1 | 1 |
| Micrococcaceae | *Arthrobacter* | 1 | 1 | 1 | 1 | 1 | 1 |
|  | *Glutamicibacter* | 0 | 0 | 2 | 2 | 0 | 0 |
|  | *Kocuria* | 4 | 4 | 6 | 6 | 2 | 2 |
|  | *Micrococcus* | 6 | 6 | 8 | 8 | 4 | 4 |
|  | *Rothia* | 1 | 1 | 0 | 0 | 0 | 0 |
| Micromonosporaceae | *Micromonospora* | 1 | 1 | 1 | 1 | 1 | 1 |
| Mycobacteriaceae | *Mycolicibacterium* | 1 | 1 | 0 | 0 | 0 | 0 |
| Nocardiaceae | *Rhodococcus* | 2 | 2 | 2 | 2 | 2 | 1 |
| Nocardoidaceae | *Aeromicrobium* | 0 | 0 | 1 | 1 | 0 | 0 |
| Nocardiopsaceae | *Nocardiopsis* | 4 | 4 | 3 | 3 | 2 | 2 |
| Ornithinimicrobiaceae | *Serinicoccus* | 1 | 1 | 0 | 0 | 0 | 0 |
| Pseudonocardiaceae | *Actinomycetospira* | 0 | 0 | 1 | 1 | 0 | 0 |
|  | *Pseudonocardia* | 1 | 1 | 0 | 0 | 0 | 0 |
| Total |  | 56 | 56 | 64 | 65 | 28 | 25 |

Six genera, namely *Gordonia*, *Jonesia*, *Mycolicibacterium*, *Pseudonocardia*, *Rothia* and *Serinicoccus* were isolated only from sponges (Table 1), which could be identified to species *Gordonia terrae* (MCC 6452), *Jonesia denitrificans* (MCC 7852), *Mycolicibacterium poriferae* (MCC 6242), *Pseudonocardia kongjuensis* (MCC 7930) and *Rothia terrae* (MCC 7823), *Serinicoccus marinus* (MCC 7935) respectively. Although 12 genera, namely *Agrococcus*, *Arthrobacter*, *Brachybacterium*, *Brevibacterium*, *Klenkia*, *Kocuria*, *Microbacterium*, *Micrococcus*, *Micromonospora*, *Nocardiopsis*, *Rhodococcus* and *Streptomyces*, were isolated from both sponges and associated habitats, most of these genera had some putative species that were either exclusive to sponges or associated environments. In particular, 7 species, *Brachybacterium muris* (MCC 7614), *Brevibacterium casei* (MCC 6140, MCC 6152, MCC 6176), *Kocuria rhizophila* (MCC 8384), *Nocardiopsis salina* (MCC 7931), *Rhodococcus zopfii* (MCC 7934), *Streptomyces smyrnaeus* (MCC 7924) and *Streptomyces viridobrunneus* (MCC 7990), were recorded only from sponges.

With respect to both, the number of isolates and number of putative species, *Streptomyces* was the most dominant genus, which was found in both sponges and associated environments. *Nocardiopsis* was the second most common genus with two dominant species *Nocardiopsis alba* (MCC 8385) followed by *N. dassonvillei* (MCC 7845). Among the genera and species that were recorded only from the environment, we provide first record of species such as *Aeromicrobium massiliense* (MCC 6739) and *Glutamicibacter mysorens* (MCC 7825) from marine waters.

### Non-obligate epibiotic predatory activity

Out of the total 50 actinobacterial isolates screened for non-obligate epibiotic predatory activity, 26 isolates showed predation on at least one of the 14 target organisms (Supplementary Table S2). Of the 26 isolates with predatory behavior, 17 preyed on Gram-negative prey, 21 preyed on Gram-positive prey, while 12 preyed on both Gram-negative and Gram-positive prey. There was no significant difference (Mann-Whitney U = 15, P = 0.2601) in the frequency of actinobacterial predators on Gram-negative and Gram-positive prey (Table 2). Most actinobacterial predators (n = 14) preyed on a single prey species while only a few predators preyed on multiple prey species. A single predator of the genus *Streptomyces* tpreyed on 8 prey species. There was a significant association between the source of isolation (sponge or associated environment) and predatory behavior (χ^2^ = 5265, P = 0.0218), where the isolates from sponge showed proportionately more predatory behavior (Figure 2).

**Figure 2.**
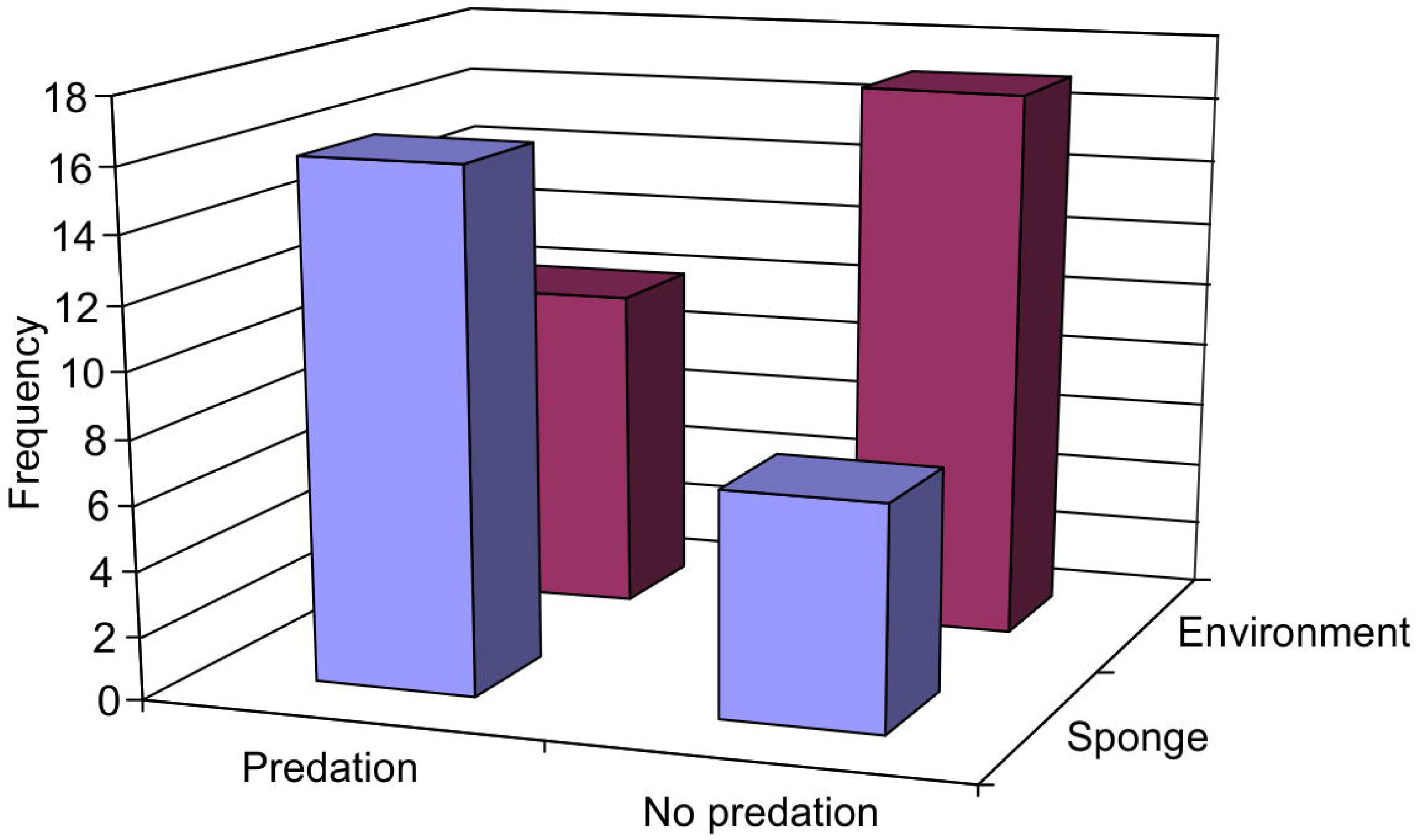
Association between source of actinobacterial isolation on their predatory behavior. There was a significant association between the source (sponge or associated environment) of actinobacterial isolation and predation (χ^2^ = 5.265, P = 0.0218).

**Table 2.** Predation and antibiotic production by actinobacteria against the Gram positive and Gram negative target species.

| Target species | Predation | Antibiotic | Predation and Antibiotic<br>by same actinobacterial<br>isolate |
| --- | --- | --- | --- |
| Gram positive |  |  |  |
| *Mycobacterium smegmatis* | 3 | 12 | 0 |
| *Micrococcus luteus* | 8 | 5 | 0 |
| *Bacillus subtilis* | 1 | 24 | 1 |
| *Staphylococcus aureus* | 17 | 9 | 4 |
| *Salinicoccus roseus* | 9 | 3 | 0 |
| *Enterococcus faecalis* | 3 | 20 | 1 |
| Gram negative |  |  |  |
| *Acetobacter pasteurianus* | 7 | 0 | 0 |
| *Alcaligenes faecalis* | 3 | 1 | 1 |
| *Escherichia coli* | 2 | 5 | 0 |
| *Klebsiella pneumoniae* | 3 | 0 | 0 |
| *Proteus vulgaris* | 8 | 0 | 0 |
| *Salmonella enterica* | 2 | 0 | 0 |
| *Serratia marcescens* | 3 | 0 | 0 |
| *Pseudomonas aeruginosa* | 1 | 0 | 0 |

All eight isolates of *Streptomyces* used for screening showed predatory behavior and preyed on both Gram-negative and Gram-positive prey (Supplementary Table S2). Out of 25 isolates of *Nocardiopsis*, 12 showed predatory behavior, out of which 5 preyed on Gram-negative bacteria while 11 preyed on Gram-positive bacteria. Both the isolates of *Micromonospora* preyed on Gram-positive prey while only one preyed on Gram-negative prey. Isolates belonging to genera *Brevibacterium*, *Glutamicibacter* and *Rhodococcus* preyed only on Gram-negative prey while *Rothia* preyed only on Gram-positive prey.

### Antibiosis, antibacterial activity and growth inhibition

Of the 50 actinobacterial isolates screened for antibacterial activity, 25 showed antibiosis against at least one target organism (Supplementary Table S2). Of these 25 isolates, all showed antibiosis against at least one of the Gram-positive target species, while only five showed antibiosis against at least one of the Gram-negative organisms. The frequency of antibacterial activity against Gram-positive organisms was significantly higher (Mann-Whitney U = 1.5, P = 0.003) than those against Gram-negative organisms (Table 2). Most antibacterial activities were broad spectrum with respect to the target organisms that they affected. There were 10 actinobacterial isolates that showed antibiosis against two target organisms, 6 isolates that affected 4 target species and 2 isolates that affected 6 target species. There was no association between antibacterial activity and the source (sponge or associated environment) of the isolation (χ^2^ = 2.0129, P = 0.1560).

Out of eight isolates of *Streptomyces* that were screened for antibacterial activity, five showed antibiosis, of which two showed antibiosis against Gram-negative target species, while all showed antibiosis against Gram-positive organisms. In the case of *Nocardiopsis*, of the 25 isolates used for screening 17 showed antibiosis, of which all affected growth of Gram-positive organisms, while only two affected growth of Gram-negative organisms. Genus *Kytococcus* showed antibiosis that affected both Gram-positive as well as Gram-negative organisms, while *Glutamicibacter* and *Rothia* showed antibiosis against Gram-positive organisms only.

### Enzyme inhibition

Out of 50 actinobacterial isolates screened for inhibition of four enzymes, 30 isolates inhibited at least one of the enzyme (Supplementary Table S2). Of these 30 isolates, 28 inhibited trypsin, 24 inhibited chymotrypsin, three inhibited angiotensin converting enzyme (ACE) and only two inhibited subtilisin. Venn diagram of frequency of isolates inhibiting different enzymes (Figure 3) suggested that five isolates inhibited only trypsin and one isolate each inhibited chymotrypsin and ACE, while subtilisin inhibition was accompanied by inhibition of other enzymes. No isolate inhibited all four enzymes. Out of 30 actinobacteria that produced enzyme inhibitors, 19 produced two inhibitors, four produced three inhibitors while seven produced only one of the four inhibitors. There was no association between the enzyme inhibition and source of the actinobacterial isolate (χ^2^ = 2.3386, P = 0.1262).

**Figure 3.**
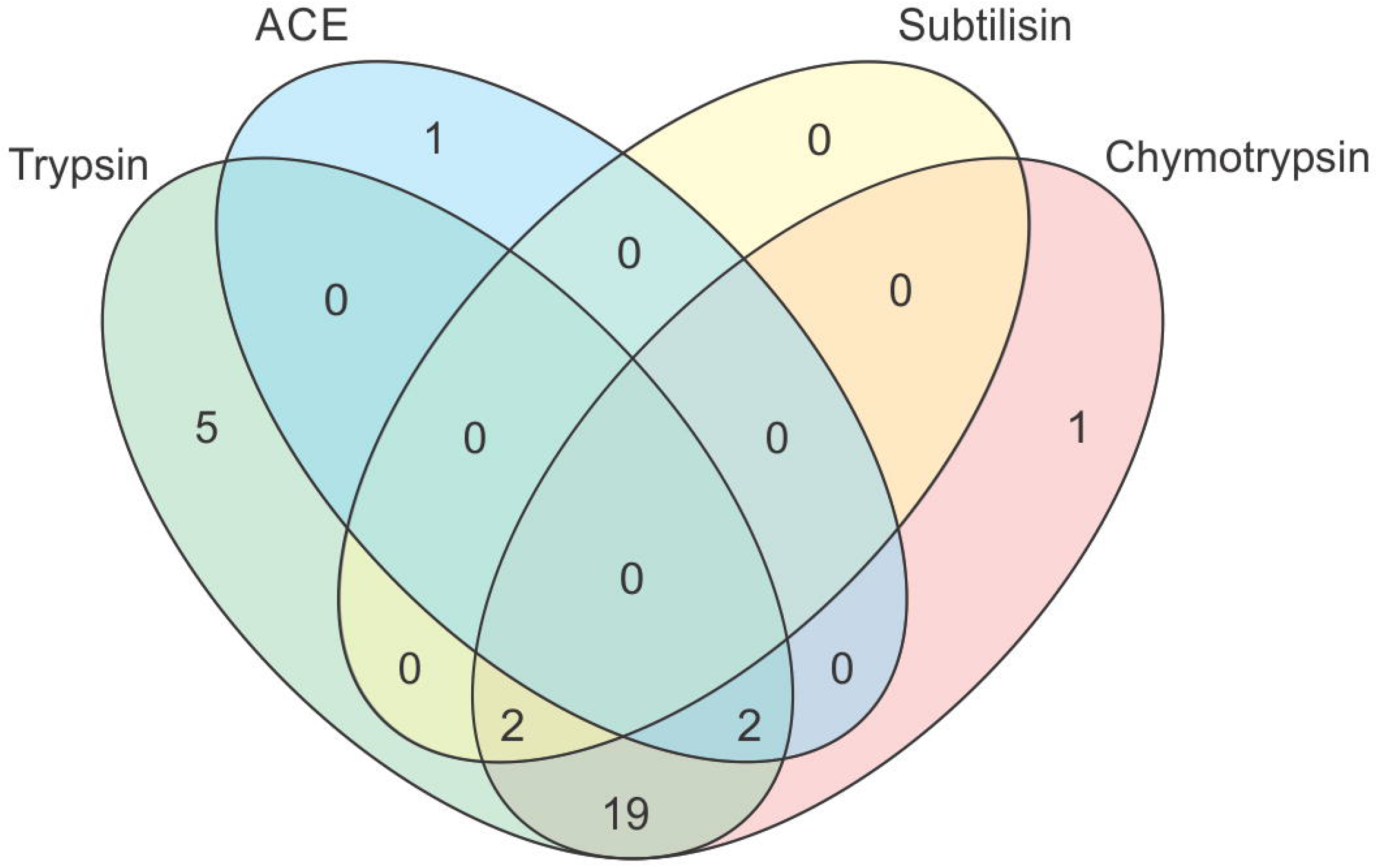
Venn diagrams combination of enzyme inhibitors produced by actinobacterial isolates. Venn diagrams is not to scale.

Out of eight isolates of *Streptomyces* seven produced enzyme inhibitors against proteases, while 12 out of 25 isolates of *Nocardiopsis* produced enzyme inhibitors of which 11 produced against proteases and two produced against ACE (Table 3). One isolate of *Actinomycetospora* inhibited activity of ACE.

**Table 3.** Frequency of actinobacterial isolates producing four different enzyme inhibitors.

| Genus | Number<br>of isolates | Frequency of isolates inhibiting |  |  |  | Isolates with<br>at least one<br>inhibition<br>activity |
| --- | --- | --- | --- | --- | --- | --- |
|  |  | Subtilisin | Trypsin | Chymotrypsin | ACE |  |
| *Actinomycetospora* | 2 | 0 | 1 | 0 | 1 | 2 |
| *Agrococcus* | 1 | 0 | 0 | 0 | 0 | 0 |
| *Brevibacterium* | 1 | 0 | 1 | 1 | 0 | 1 |
| *Glutamicibacter* | 1 | 0 | 1 | 1 | 0 | 1 |
| *Jonesia* | 1 | 0 | 0 | 0 | 0 | 0 |
| *Kocuria* | 1 | 0 | 0 | 0 | 0 | 0 |
| *Kytococcus* | 1 | 0 | 1 | 0 | 0 | 1 |
| *Micrococcus* | 1 | 0 | 1 | 0 | 0 | 1 |
| *Micromonospora* | 2 | 0 | 2 | 2 | 0 | 2 |
| *Nocardiopsis* | 25 | 0 | 11 | 11 | 2 | 12 |
| *Pseudonocardia* | 1 | 0 | 0 | 0 | 0 | 0 |
| *Rhodococcus* | 4 | 0 | 2 | 1 | 0 | 2 |
| *Rothia* | 1 | 0 | 1 | 1 | 0 | 1 |
| *Streptomyces* | 8 | 2 | 7 | 7 | 0 | 7 |

### Associations between different activities

Out of 50 actinobacterial isolates that were screened for activities, 39 showed at least one of the three activities. Of these 39 isolates, 15 showed all three activities, while nine showed predation as well as enzyme inhibition (Figure 4). There were only seven isolates that showed predation and antibiotic production against the same target organism (Table 2) and all these isolates belonged to genera *Streptomyces* and *Nocardiopsis*.

**Figure 4.**
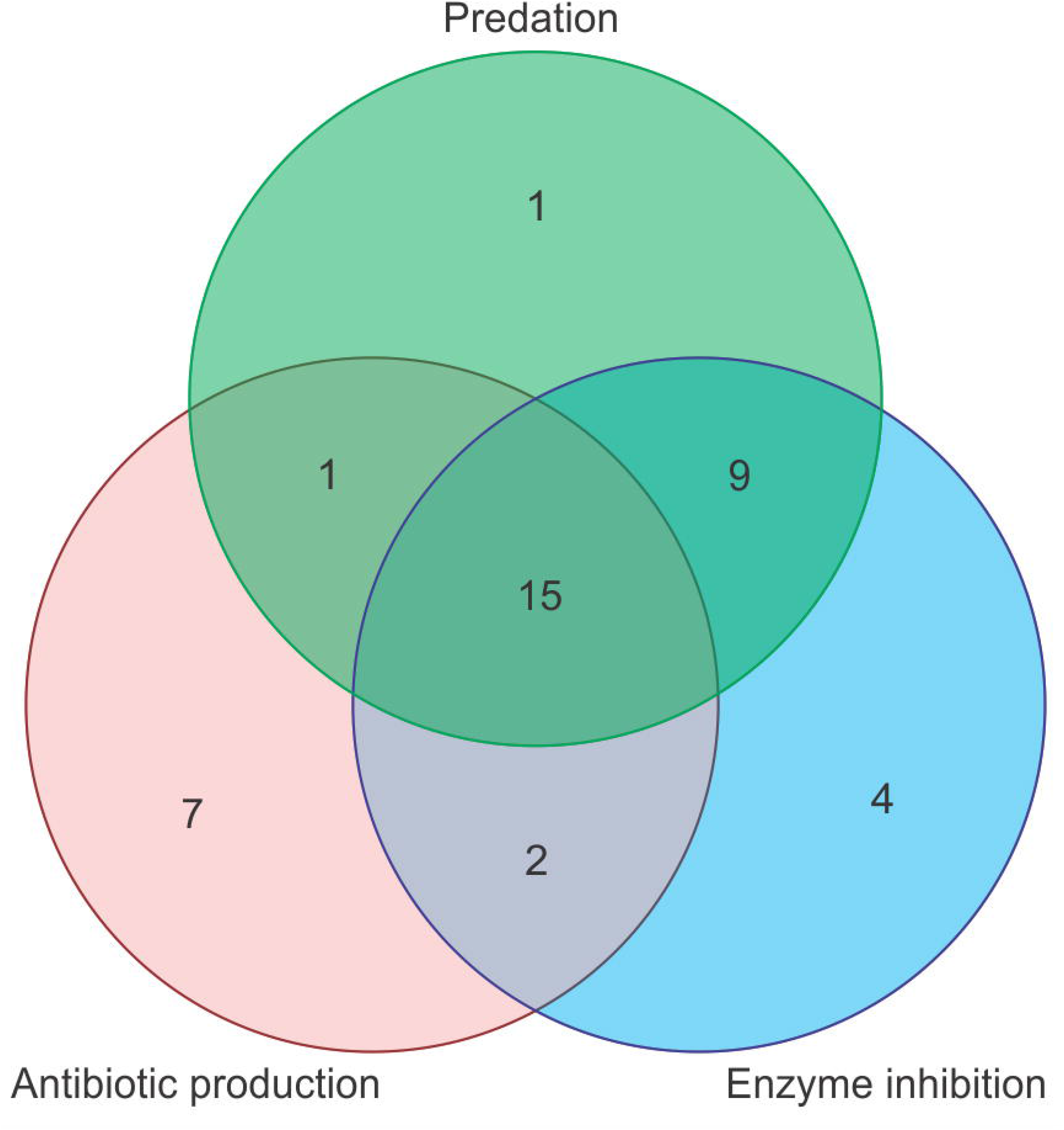
Venn diagrams of predation, antibiotic production and enzyme inhibition by actinobacterial isolates. Venn diagrams is not to scale.

Antibiotic production showed no significant association with predation (χ^2^ = 2.8846, P = 0.0894) or any of the four enzyme inhibition (χ^2^ = 2.0525, P = 0.1520). However, there were significant associations between predation and protease inhibitors (Figure 5). There were 24 isolates that showed both predation as well as inhibition of at least one enzyme and there was a significant association between the two activities (χ^2^ = 26.172, P < 0.0001), where predators proportionately produced more enzyme inhibitors than non-predators (Figure 5a). There were 23 actinobacterial isolates that showed predation as well as trypsin inhibition and there was a significant association between the two (χ^2^ = 23.165, P < 0.0001) with predators more likely to produce trypsin inhibitors than non-predators (Figure 5b). Similarly, 24 actinobacteria were predators as well as inhibited chymotrypsin activity and there was a significant association between the two (χ^2^ = 42.604, P < 0.0001) with predators more likely to produce chymotrypsin inhibitors than non-predators (Figure 5c).

**Figure 5.**
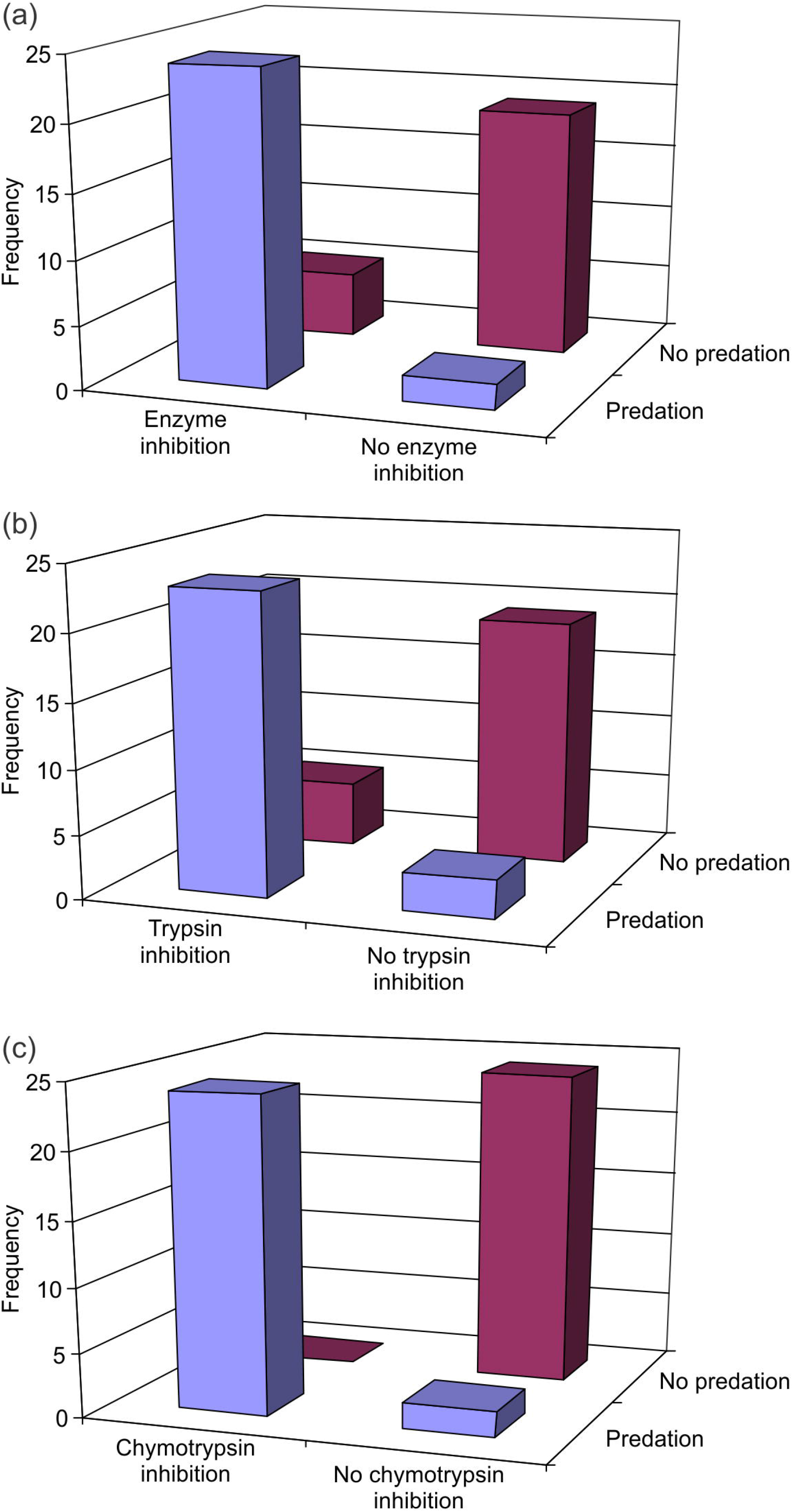
Association between enzyme inhibition and predation in actinobacterial isolates. Predation was significantly associated with (a) inhibition of any one of the four enzymes tested (χ^2^ = 26.172, P < 0.0001), (b) inhibition of trypsin (χ^2^ = 23.165, P < 0.0001) and (c) inhibition of chymotrypsin (χ^2^ = 42.604, P < 0.0001).

## DISCUSSION

Sponges and associated environment in northern parts of western coast of India are rich in actinobacterial diversity with about 95 putative species under 19 families and 28 genera. We recorded 13 species of actinobacteria only from sponges. Out of these, *Mycobacterium poriferae* was originally described from marine sponge (Padgitt and Moshier, 1987), while three species, *Gordonia terrae* (Elfalah et al., 2013; Santos et al., 2019; Montalvo et al., 2005), *Brevibacterium casei* (Kiran et al., 2010) and *Koc uria rhizophila* (Palomo et al., 2013), have been previously reported from sponges. To our knowledge, we provide first report of nine species, namely *Brachybacterium murisi, Jonesia denitrificans*, *Nocardiopsis salina*, *Pseudonocardia kongjuensis*, *Rhodococcus zopfii*, *Rothia terrae*, *Serinicoccus marinus*, *Streptomyces smyrnaeus* and *Streptomyces viridobrunneus*, from marine sponges, although some of them are known from marine habitats (Stach et al., 2003; Satheeja and Jebakumar, 2011; Yi et al., 2004; Shinde et al., 2018).

*Streptomyces* was the most dominant genus among the isolates, which agrees with the findings of Zhang et al. (2008). Genus *Nocardiposis*, with its two species *N. alba* and *N. dassonvillei*, has been suggested (Bennur et al., 2015) as the second common genus after *Streptomyces* and that too agrees with our findings. Further, report of most genera, including *Agrococcus*, *Arthrobacter*, *Brevibacterium*, *Kocuria*, *Microbacterium* and *Micrococcus*, from sponges in our study are consistent with previous reports from other study areas including South China Sea (Li et al., 2015), Yellow Sea (Zhang et al., 2008), Mediterranean Sea (Cheng et al., 2015), coast of Florida in USA (Montalvo, 2005) and northern coast of Brazil (Menezes et al., 2010) indicating that there are common trends in the discovery of actinobacteria from sponges.

Among the first reports from marine environment from our study, *Aeromicrobium massiliense* and *Glutamicibacter mysorens* are known from human fecal microbiota (Ramasamy et al., 2012) and sewage (Nand and Rao, 1972) respectively. Presence of these two species in the sediments along the collection site Harne (17.81°N, 73.09°E) likely suggests fecal pollution in this area.

Although predation is a widespread behavior in bacterial kingdom, δ-proteobacteria of the orders *Myxococcales* and *Bdellovibrionales* have received more attention (Jurkevitch, 2007) as compared to other taxa, especially the Gram-positive bacteria such as actinobacteria. Among actinobacteria only three genera, namely *Agromyces*, *Streptomyces* and *Streptoverticillium*, are known to have predatory behavior against other bacterial species (Casida, 1980; 1983; 1988; Kumbhar et al., 2014; Zeph and Casida, 1986; Ibrahimi et al., 2020). In the current study, for the first time, we show predation in six other genera of actinobacteria, namely *Brevibacterium*, *Glutamicibacter*, *Micromonospora*, *Nocardiopsis*, *Rhodococcus* and *Rothia*. Kumbhar et al. (2014) argued that predatory behavior is widespread in genus *Streptomyces* and even in the current study we observed that all the isolates of *Streptomyces* used for screening showed predation on Gram-positive as well as Gram-negative prey.

Since sponges are sessile and lack other anti-predator defenses, it has been suggested that secondary metabolites of bacteria can provide sponges with chemical defense (Lee et al., 2001; Kumbhar and Watve, 2013). However, we did not observe any significant association between the source of actinobacterial isolation and antibiotic production, suggesting that isolates even from environment were equally likely to produce antimicrobials as that of the isolates recovered from sponges. However, there was a significant association between the source of isolation and predatory activity, with proportionately more predators among the isolates recovered from sponge. Ecologically this makes sense. As the sponges are filter feeders and have regular intake of environmental bacteria, sponge associated actinobacteria will have better predation opportunities. It is also possible that the predatory activity of sponge associated actinobacteria, could have evolved as a mutualistic activity as it can defend sponges from pathogenic bacterial invasions.

Actinobacteria are known to produce several enzyme inhibitors (Manivasagan et al., 2015; Imada, 2005). However, for the first time we show a strong association between predation and enzyme inhibition, specifically inhibition of trypsin and chymotrypsin, where predators produced proportionality more enzyme inhibitors as compared to non predators. Predators themselves are known to produce a variety of hydrolytic enzymes for degrading the prey (Pérez et al., 2016). Therefore, it is possible that the production of enzyme inhibitors safeguards their own cells from being target of the enzyme. It is also possible that enzyme inhibitors also protect the actinobacteria from hydrolytic enzymes produced from the sponge host and other microbiota.

An interesting observation that we made, when comparing the predation and antibiotic production by actinobacteria, was that, while predation was equally effective against Gram-positive as well as Gram-negative target species, antibiotic production was mainly effective against Gram-positive bacteria. Recently, Ibrahimi et al. (2020) suggested that there are some bio-active secondary metabolites that co-cultured actinobacteria produce in the presence of prey cells. It is therefore possible that studying the predatory behavior of actinobacteria and predation specific metabolites could lead to discovery of novel therapeutic agents that are more broad-spectrum.

Although actinobacteria are known to be rich in secondary metabolites, extracellular enzymes and enzyme inhibitors, the ecological role of these extracellular bioactive molecules is little known. We suggest that studying the ecological correlates of bioactivity and the inter-correlation patterns of different types of bioactivity can be a useful tool in understanding the ecological origins of bioactivity and testing alternative ecological hypotheses.

## CONCLUSION

Sponges and associated environments of intertidal zones, along the northern parts of west coast of India, are rich in actinobacterial diversity with 19 families and 28 genera, which could be attributed to 95 putative species using mPTP and 100 putative species based on bPTP methods. Although, at the genus level, the trends in the discovery of actinobacteria isolated from sponges was consistent with previous studies from different study areas, we provide first report of nine species, namely *Brachybacterium murisi, Jonesia denitrificans*, *Nocardiopsis salina*, *Pseudonocardia kongjuensis*, *Rhodococcus zopfii*, *Rothia terrae*, *Serinicoccus marinus*, *Streptomyces smyrnaeus* and *Streptomyces viridobrunneus*. Non-obligate epibiotic predatory behavior was widespread among actinobacterial genera and we provide first report of predatory activity in *Brevibacterium*, *Glutamicibacter*, *Micromonospora, Nocardiopsis, Rhodococcus* and *Rothia*. Sponges associated actinobacteria showed significantly more predatory behavior than environmental isolates, and we hypothesize that predatory actinobacteria might provide sponges with defense against pathogenic bacteria. While antibiotic produced from actinobacterial isolates affected Gram-positive target bacteria with little to no effect on Gram-negative bacteria, predation targeted both Gram-positive and Gram-negative prey with equal propensity, suggesting that study of predation specific metabolites might provide novel therapeutic agents with broad-spectrum. Actinobacterial isolates from both sponge and associated environment produced inhibitors of serine proteases and angiotensin converting enzyme. Predatory behavior was strongly associated with inhibition of trypsin and chymotrypsin, which might be helpful for the actinobacteria for overcoming effects of proteolytic enzymes produced by sponge host and other microbiota. Understanding diversity and associations among various actinobacterial activities, with each other and the source of isolation, can provide new insights in marine microbial ecology and provide opportunities to isolate novel therapeutic agents.

## DATA AVAILABILITY

Sequences of 16S rRNA gene of studied isolates are submitted to GenBank NCBI under the accession numbers MN339687–MN339897 and MT598037–MT598065. Actinobacterial cultures are deposited in the Microbial Culture Collection (MCC) of National Centre for Microbial Resource, National Center for Cell Sciences, Pune, India (accession numbers are provided in the Supplementary Table S1). All the data used for analysis is provided in supplementary information (Supplementary Table S1 and Table S2).

## ACKNOWLEDGEMENTS

This work was funded by Maharashtra Gene Bank Programme (RGSTC/File-2007/DPP-054/CR-28) of Rajiv Gandhi Science and Technology Commission, Government of Maharashtra, India. We thank the Director and Chair of Biology, Indian Institute of Science Education and Research (IISER), Pune, and Principle, M.E.S. Abasaheb Garware College, Pune for providing infrastructural facilities. We are thankful to V. S. Rao, IISER Pune, for support an encouragement.

## AUTHOR CONTRIBUTIONS

M.W., U.B. and N. Deshpande conceived and designed the study. U.B., N.S., K.H., A.P., U.L., T.G., K.P., A.J., R.S., H.V and V.T. performed the study. N. Dahanukar and M.W. analyzed the data. N. Dahanukar, U.B. and M.W. wrote the manuscript with inputs from other authors. All authors contributed to the proofreading of the manuscript.

## Supplementary information

Supplementary Table S1 and Table S2 accompanies the online version of the paper.

## References

Abdelfattah, M. S., Elmallah, M. I. Y., Hawas, U. W., El-Kassema, L. T. A., and Eid, M. A. G. (2016). Isolation and characterization of marine-derived actinomycetes with cytotoxic activity from the Red Sea coast. Asian Pacific J. Tropical Biomed. 6, 651–657.

Altschul, S. F., Gish, W., Miller, W., Myers, E. W., and Lipman, D. J. (1990). Basic local alignment search tool. J. Mol. Biol. 215, 403–410.

Barka, E. A., Vatsa, P., Sanchez, L., Gaveau-Vaillant, N., Jacquard, C., Meier-Kolthoff, J. P., Klenk, H. P., Clément, C., Ouhdouch, Y., and van Wezel, G. P. (2016). Taxonomy, physiology, and natural products of Actinobacteria. Microbiol. Mol. Biol. Rev. 80, 1–43.

Bennur, T., Kumar, A. R., Zinjarde, S., and Javdekar, V. (2015). *Nocardiopsis* species: Incidence, ecological roles and adaptations. Microbiol. Res. 174, 33–47.

Braesel, J., Lee, J. H., Arnould, B., Murphy, B. T., and Eustáquio, A. S. (2019). Diazaquinomycin biosynthetic gene clusters from marine and freshwater actinomycetes. J. Nat. Prod. 82, 937–946.

Casida, L. E. Jr. (1980). Bacterial predators of *Micrococcus luteus* in soil. Appl. Environ. Microbiol. 39, 1035–1041.

Casida, L. E. Jr. (1983). Interaction of *Agromyces ramosus* with other bacteria in soil. Appl. Environ. Microbiol. 46, 881–888.

Casida, L. E. (1988). Minireview: Nonobligate bacterial predation of bacteria in soil. Microb. Ecol. 15, 1–8.

Cheng, C., MacIntyre, L., Abdelmohsen, U. R., Horn, H., Polymenakou, P. N., Edrada-Ebel, R., and Hentschel, U. (2015). Biodiversity, anti-trypanosomal activity screening, and metabolomic profiling of actinomycetes isolated from Mediterranean sponges. PLoS ONE 10, e0138528. doi: 10.1371/journal.pone.0138528.

Cheung, A. L., Ying, P., and Fischetti, V. A. (1991). A method to detect proteinase activity using unprocessed X-ray films. Anal. Biochem. 193, 20–23.

Cushman, D. W., and Cheung, H. S. (1971). Spectrophotometric assay and properties of the angiotensin-converting enzyme of rabbit lung. Biochem. Pharmacol. 20, 1637–1648.

Edgar, R. C. (2004). MUSCLE: multiple sequence alignment with high accuracy and high throughput. Nucleic Acids Res. 32, 1792–1797.

Elfalah, H. W., Usup, G., and Ahmad, A. (2013). Anti-microbial properties of secondary metabolites of marine *Gordonia tearrae* extract. J. Agric. Sci. 5, 94–101.

Gandhimathi, R. Arunkumar, M., Selvin, J., Thangavelu, T., Sivaramakrishnan, S., Kiran, G. S., Shanmughapriya, S., and Natarajaseenivasana, K. (2008). Antimicrobial potential of sponge associated marine actinomycetes. J. Mycol. Med. 18, 16–22.

Goodfellow, M. (2015) “Actinobacteria phyl. nov.”, in Bergey’s Manual of Systematics of Archaea and Bacteria, eds Whitman, W. B. et al., (New York: Wiley), 1–2. doi: 10.1002/9781118960608.pbm00002.

Harir, M., Bendif, H., Bellahcene, M., Fortas, Z. & Pogni, R. (2018). *“Streptomyces* secondary metabolites”, in Basic Biology and Applications of Actinobacteria, ed. Enany, S. (London, IntechOpen), 99–122.

Hoang, D. T., Chernomor, O., von Haeseler, A., Minh, B. Q., and Vinh, L.S. (2018). UFBoot2: Improving the ultrafast bootstrap approximation. Mol. Biol. Evol. 35, 518–522.

Ibrahimi, M., Korichi, W., Hafidi, M., Lemee, L., Ouhdouch, Y., and Loqman, S. (2020). Marine Actinobacteria: Screening for Predation Leads to the Discovery of Potential New Drugs against Multidrug-Resistant Bacteria. Antibiotics 9(2), 91.

Imada, C. (2005). Enzyme inhibitors and other bioactive compounds from marine actinomycetes. Antonie Van Leeuwenhoek 87, 59–63.

Jose, P. A., and Jebakumar, S. R. D. (2014). Unexplored hypersaline habitats are sources of novel actinomycetes. Front. Microbiol. 5, 242. doi: 10.3389/fmicb.2014.00242.

Jurkevitch, E. (2007). Predatory behaviors in bacteria-diversity and transitions. Microbe 2, 67–73.

Kalyaanamoorthy, S., Minh, B. Q., Wong, T. K. F., von Haeseler, A., and Jermiin, L.S. (2017). ModelFinder: Fast model selection for accurate phylogenetic estimates. Nat. Methods 14, 587–589.

Kiran, G. S., Sabarathnam, B., and Selvin, J. (2010). Biofilm disruption potential of a glycolipid biosurfactant from marine *Brevibacterium casei*. FEMS Immunol. Med. Microbiol. 59, 432–438.

Kokare, C. R., Mahadik, K. R., and Kadam, S. S. (2004). Isolation of bioactive marine actinomycetes from sediments isolated from Goa and Maharashtra coastlines (west coast of India). Indian J. Mar. Sci. 33, 248–256.

Kumar, S., Stecher, G., and Tamura, K. (2016). MEGA7: molecular evolutionary genetics analysis version 7.0 for bigger datasets. Mol. Biol. Evol. 33, 1870–1874.

Kumbhar, C., and Watve, M. (2013). Why antibiotics: A comparative evaluation of different hypotheses for the natural role of antibiotics and an evolutionary synthesis. Nat. Sci. 5, 26–40.

Kumbhar, C., Mudliar, P., Bhatia, L., Kshirsagar, A., and Watve, M. (2014). Widespread predatory abilities in the genus *Streptomyces*. Arch. Microbiol. 196, 235–248.

Lam, K. S. (2006). Discovery of novel metabolites from marine actinomycetes. Curr. Opin. Microbiol. 9, 245–251.

Lee, O. O., Wong, Y. H., and Qian, P. Y. (2009). Inter- and intraspecific variations of bacterial communities associated with marine sponges from San Juan Island, Washington. Appl. Environ. Microbiol. 75, 3513–3521.

Lee, Y. K., Lee, J. H., and Lee, H. K. (2001). Microbial symbiosis in marine sponges. J. Microbiol. 39, 254–264.

Li, Z., Sun, W., Zhang, F., He, L., and Loganathan, K. (2015). Actinomycetes from the South China Sea sponges: isolation, diversity and potential for aromatic polyketides discovery. Front. Microbiol. 6, 1048. doi: 10.3389/fmicb.2015.01048.

Mahapatra, G. P., Raman, S., Nayak, S., Gouda, S., Das, G., and Patra, J. K. (in press). Metagenomics approaches in discovery and development of new bioactive compounds from marine actinomycetes. Curr. Microbiol. doi: 10.1007/s00284-019-01698-5.

Mahmoud, H. M., and Kalendar, A. A. (2016) Coral-associated Actinobacteria: diversity, abundance, and biotechnological potentials. Front. Microbiol. 7, 204. doi: 10.3389/fmicb.2016.00204.

Manivasagan, P., Kang, K. H., Sivakumar, K., Li-Chan, E. C., Oh, H. M., and Kim, S. K. (2014). Marine actinobacteria: an important source of bioactive natural products. Environ. Toxicol. Pharmacol. 38, 172–188.

Manivasagan, P., Venkatesan, J., Sivakumar, K., and Kim, S. K. (2015). Actinobacterial enzyme inhibitors–A review. Crit. Rev. Microbiol. 41, 261–272.

Menezes, C. B., Bonugli-Santos, R. C., Miqueletto, P. B., Passarini, M. R., Silva, C. H., Justo, M. R., Leal, R. R., Fantinatti-Garboggini, F., Oliveira, V. M., Berlinck, R. G., and Sette, L. D. (2010). Microbial diversity associated with algae, ascidians and sponges from the north coast of São Paulo state, Brazil. Microbiol. Res. 165, 466–482.

Mincer, T. J., Jensen, P. R., Kauffman, C. A., and Fenical, W. (2002). Widespread and persistent populations of a major new marine actinomycete taxon in ocean sediments. Appl. Environ. Microbiol. 68, 5005–5011.

Mohammadipanah, F., and Wink, J. (2016). Actinobacteria from arid and desert habitats: diversity and biological activity. Front. Microbiol. 6, 1541. doi: 10.3389/fmicb.2015.01541.

Montalvo, N. F., Mohamed, N. M., Enticknap, J. J., and Hill, R. T. (2005). Novel actinobacteria from marine sponges. Antonie Van Leeuwenhoek 87, 29–36.

Nand, K., and Rao, D. V. (1972). *Arthrobacter mysorens–a* new species excreting L-glutamic acid. Zentralbl Bakteriol Parasitenkd Infektionskr Hyg. 127, 324–331.

Nei, M., and Kumar, S. (2000). Molecular evolution and phylogenetics. UK, Oxford University Press.

Ng, K.H., Lye, H.S., Easa, A.M., and Liong, M.T. (2008). Growth characteristics and bioactivity of probiotics in tofu-based medium during storage. Ann. Microbiol. 58, 477–487.

Nguyen, L. T., Schmidt, H. A., von Haeseler, A., and Minh, B. Q. (2015). IQ-TREE: a fast and e ective stochastic algorithm for estimating maximum likelihood phylogenies. Mol. Biol. Evol. 32, 268–274.

Olano, C., Méndez, C., and Salas, J. (2009). Antitumor compounds from marine actinomycetes. Marine Drugs 7, 210–248.

Padgitt, P. J., and Moshier, S. E. (1987). *Mycobacterium poriferae* sp. nov., a scotochromogenic, rapidly growing species isolated from a marine sponge. Int. J. Syst. Evol. Microbiol. 37, 186–191.

Palomo, S., González, I, de la Cruz, M., Martín, J., Tormo, J. R., Anderson, M., Hill, R. T., Vicente, F., Reyes, F., and Genilloud1, O. (2013). Sponge-derived *Kocuria* and *Micrococcus* spp. as sources of the new thiazolyl peptide antibiotic kocurin. Marine Drugs 11, 1071–1086.

Pathom-Aree, W., Stach, J. E., Ward, A. C., Horikoshi, K., Bull, A. T., and Goodfellow, M. (2006). Diversity of actinomycetes isolated from Challenger Deep sediment (10,898 m) from the Mariana Trench. Extremophiles 10, 181–189.

Pérez, J., Moraleda-Muñoz, A., Marcos-Torres, F. J., and Muñoz-Dorado, J. (2016). Bacterial predation: 75 years and counting! Environ. Microbiol. 18, 766–779.

Pimentel-Elardo, S. M., Kozytska, S., Bugni, T. S., Ireland, C. M., Moll, H., and Hentschel, U. (2010). Anti-parasitic compounds from *Streptomyces* sp. strains isolated from Mediterranean sponges. Marine Drugs 8, 373–380.

Ramasamy, D., Kokcha, S., Lagier, J.-C., Nguyen, T.-T., Raoult, D., and Fournier, P.-E. (2012). Genome sequence and description of *Aeromicrobium massiliense* sp. nov. Stand. Genomic Sci. 7, 246–257.

Rambaut, A. (2009). FigTree. ver 1.4.3 http://tree.bio.ed.ac.uk/software/%ef%ac%81gtree/.

Riquelme, C., Hathaway, J. J. M., Enes Dapkevicius, M. de L. N., Miller, A. Z., Kooser, A., Northup, D. E., Jurado, V., Fernandez, O., Saiz-Jimenez, C., and Cheeptham, N., (2015). Actinobacterial diversity in volcanic caves and associated geomicrobiological interactions. Front. Microbiol. 6, 1342. doi: 10.3389/fmicb.2015.01342.

Santos, J. D., Vitorino, I., De la Cruz, M., Díaz, C., Cautain, B., Annang, F., Pérez-Moreno, G., Martinez, I. G., Tormo, J. R., Martín, J. M., Urbatzka, R., Vicente, F. M., and Lage, O. M. (2019). Bioactivities and extract dereplication of Actinomycetales isolated from marine sponges. Front. Microbiol. 10, 727. doi: 10.3389/fmicb.2019.00727.

Satheeja, S. V., and Jebakumar, S. R. (2011). Phylogenetic analysis and antimicrobial activities of *Streptomyces* isolates from mangrove sediment. J. Basic Microbiol. 51, 71–79.

Schwarz, G. (1978). Estimating the dimension of a model. Ann. Stat. 6, 461–464.

Shinde, V.L., Meena, R.M., and Shenoy, B.D., (2018). Phylogenetic characterization of culturable bacteria and fungi associated with tarballs from Betul beach, Goa, India. Mar. Pollut. Bull. 128, 593–600.

Shivlata, L., and Tulasi, S. (2015). Thermophilic and alkaliphilic Actinobacteria: biology and potential applications. Front. Microbiol. 6, 1014. doi: 10.3389/fmicb.2015.01014.

Stach, J. E. M., Maldonado, L. A., Masson, D. G., Ward, A. C., Goodfellow, M., Bull, A. T. (2003). Statistical approaches for estimating actinobacterial diversity in marine sediments. Appl. Environ. Microbiol. 69, 6189–6200.

Subramani, R., and Sipkema, D. (2019). Marine rare actinomycetes: A promising source of structurally diverse and unique novel natural products. Marine Drugs 17, 249.

Tan, L. T. H., Ser, H. L., Yin, W. F., Chan, K. G., Lee, L. H., and Goh, B. H. (2015). Investigation of antioxidative and anticancer potentials of *Streptomyces* sp. MUM256 isolated from Malaysia mangrove soil. Front. Microbiol. 6, 1316. doi: 10.3389/fmicb.2015.01316.

Taylor, M. W., Radax, R., Steger, D., and Wagner, M. (2007). Sponge-associated microorganisms: evolution, ecology, and biotechnological potential. Microbiol. Mol. Biol. Rev. 71, 295–347.

Thomas, T. R., Kavlekar, D. P., and LokaBharathi, P. A. (2010). Marine drugs from sponge-microbe association - a review. Marine Drugs 8, 1417–1468.

Tripathi, V. R., Kumar, S., and Garg, S. K. (2011). A study on trypsin, *Aspergillus flavus* and *Bacillus* sp. protease inhibitory activity in *Cassia tora* (L.) syn *Senna tora* (L.) Roxb. seed extract. BMC Complement. Altern. Med. 11, 56. doi: 10.1186/1472-6882-11-56.

Trischman, J. A., Tapiolas, D. M., Jensen, P. R., Dwight, R., Fenical, W., McKee, T. C., Ireland, C. M., Stout, T. J., and Clardy, J. (1994). Salinamides A and B: anti-inflammatory depsipeptides from a marine streptomycete. J. American Chem. Soc. 116, 757–758.

Trujillo, M. E., Riesco, R., Benito, P., and Carro, L. (2015). Endophytic actinobacteria and the interaction of *Micromonospora* and nitrogen fixing plants. Front. Microbiol. 6, 1341. doi: 10.3389/fmicb.2015.01341.

Valli, S., Suvathi, S. S., Aysha, O. S., Nirmala, P., Vinoth, K. P., and Reena, A. (2012). Antimicrobial potential of Actinomycetes species isolated from marine environment. Asian Pacific J. Tropical Biomed. 2, 469–473.

Van der Meij, A., Worsley, S. F., Hutchings, M. I., and van Wezel, G. P. (2017). Chemical ecology of antibiotic production by actinomycetes. FEMS Microbiol. Rev. 41, 392–416.

Velho-Pereira, S., and Kamat, N.M. (2011). Antimicrobial screening of actinobacteria using a modified cross-streak method. Indian J. Pharm. Sci. 73, 223–228.

Ward, A. C., and Bora, N. (2006). Diversity and biogeography of marine actinobacteria. Curr. Opin. Microbiol. 9, 279–286.

Watve, M., Shejval, V., Sonawane, C., Rahalkar, M., Matapurkar, A., Shouche, Y., Patole, M., Phadnis, N., Champhenkar, A., Damle, K., Karandikar, S., Kshirsagar, V., and Jog, M. (2000). The ‘K’ selected oligophilic bacteria: A key to uncultured diversity? Curr. Sci. 78, 1535–1542.

Watve, M. G., Tickoo, R., Jog, M. M., and Bhole, B. D. (2001) How many antibiotics are produced by the genus Streptomyces? Arch. Microbiol. 176, 386–390.

Yang, J., Li, X., Huang, L., and Jiang, H. (2015). Actinobacterial diversity in the sediments of five cold springs on the qinghai-tibet plateau. Front. Microbiol. 6, 1345. doi: 10.3389/fmicb.2015.01345.

Yi, H., Schumann, P., Sohn, K. and Chun, J. (2004). Serinicoccus marinus gen. nov., sp. nov., a novel actinomycete with L-ornithine and L-serine in the peptidoglycan. Int. J. Syst. Evol. Microbiol. 54(5), 1585–1589.

Zeph, L. R., and Casida, L. E. (1986). Gram-negative versus gram-positive (actinomycete) nonobligate bacterial predators of bacteria in soil. Appl. Environ. Microbiol. 52, 819–823.

Zhang, H., Zhang, W., Jin, Y., Jin, M., and Yu, X. (2008). A comparative study on the phylogenetic diversity of culturable actinobacteria isolated from five marine sponge species. Antonie van leeuwenhoek 93, 241–248.

Zhang, J., Kapli, P., Pavlidis, P., and Stamatakis, A. (2013). A general species delimitation method with applications to phylogenetic placements. Bioinformatics 29, 2869–2876.

ZoBell, C. E. (1941). Studies on marine bacteria. I. The cultural requirements of heterotrophic aerobes. J. Mar. Res. 4, 41–75.

